# Improving Mathematical Models of Cancer through Game-Theoretic Modelling: A Study in Non-Small Cell Lung Cancer

**DOI:** 10.1101/2021.10.29.466444

**Authors:** Virginia Ardévol Martínez, Monica Salvioli, Narmin Ghaffari Laleh, Frank Thuijsman, Joel S. Brown, Rachel Cavill, Jakob Nikolas Kather, Kateřina Staňková

**Affiliations:** Department of Data Science and Knowledge Engineering, Maastricht University, Maastricht, The Netherlands; Faculty of Technology, Policy and Management, Delft University of Technology, Delft, The Netherlands; Department of Medicine III, University Hospital RWTH Aachen, Aachen, Germany; Department of Integrated Mathematical Oncology, Moffitt Cancer Center, Tampa, Florida, United States of America; Medical Oncology, National Center for Tumor Diseases, University Hospital Heidelberg, Heidelberg, Germany

**Keywords:** Evolutionary game theory, Non-Small Cell Lung Cancer, immunotherapy, radiotherapy, Stackelberg evolutionary game, therapy resistance

## Abstract

We examined a dataset of 590 Non-Small Cell Lung Cancer patients treated with either chemotherapy or immunotherapy using a game-theoretic model that includes both the evolution of therapy resistance and a cost of resistance. We tested whether the game-theoretic model provides a better fit than classical mathematical models of population growth (exponential, logistic, classic Bertalanffy, general Bertalanffy, Gompertz, general Gompertz). To our knowledge, this is the first time a large clinical patient cohort (as opposed to only in-vitro data) has been used to apply a game-theoretic cancer model. The game-theoretic model provided a better fit to the tumor dynamics of the 590 Non-Small Cell Lung Cancer patients than any of the non-evolutionary population growth models. This was not simply due to having more parameters in the game-theoretic model. The game-theoretic model was seemingly able to fit more accurately patients whose tumor burden exhibit a U-shaped trajectory over time. We explained how this game-theoretic model provides predictions of future tumor growth based on just a few initial measurements. Using the estimates for treatment-specific parameters, we then explored alternative treatment protocols and their expected impact on tumor growth and patient outcome. As such, the model could possibly be used to suggest patient-specific optimal treatment regimens with the goal of minimizing final tumor burden. Therapeutic protocols based on game-theoretic modeling can help to predict tumor growth, and could potentially improve patient outcome in the future. The model invites evolutionary therapies that anticipate and steer the evolution of therapy resistance.

## 1 Introduction

Lung cancer is the second most common cancer and the leading cause of cancer death for both men and women and Non-Small Cell Lung Cancer (NSCLC) is the most frequent type of lung cancer, accounting for 84% of all lung cancer diagnoses [7]. Although novel anti-cancer therapies like targeted therapies and immunotherapy are allowing people with metastatic lung cancer to live longer than ever before, metastatic lung cancer remains incurable [7, 5]. This is often caused by the evolution of therapy resistance that the therapy selects for [23, 50, 40, 30].

To improve therapy, we need to better understand tumor response to different potential therapy options [**?**]. Mathematical modelling helps with this understanding [2]. Here we compare the fits of six mathematical models (exponential, logistic, classic Bertalanffy, general Bertalanffy, Gompertz and general Gompertz) with that of a novel game-theoretic model. The data were from patients with Non-small Cell Lung Cancer from four clinical trials (NCT01846416 GO28625 FIR, NCT01903993 GO28753 POPLAR, NCT02031458 GO28754 BIRCH, NCT02008227 GO28915 OAK) [24]. For each patient there were measurements of the diameter of a patient’s tumor taken over time. Recently, Ghaffari Laleh et al. (2022) had fitted classic models of tumor growth to this dataset and performed a comparison between them, showing that they can fit the trajectory of tumor growth relatively well [24]. They can provide a good estimation of the future response based on early treatment data. However, the models’ predictive capabilities fail when tumor growth is not monotonic, i.e., for tumors that are neither continuously growing nor shrinking [24]. Yet, many tumor growth dynamics show a U-shape where therapy is initially effective but loses efficacy as the cancer cell population evolves resistance.

In this paper, we use a subset of 590 patients from Ghaffari Laleh et al. (2022), corresponding to those patients with NSCLC treated with chemotherapy or immunotherapy for which we have measurements for at least 6 time points. We propose a game-theoretic model which assumes that evolution of therapy resistance is a quantitative trait. This is different from some existing models that assume a qualitative trait where different cancer cell types are either resistant or sensitive to treatment [58, 33, 5]. The model we utilize here has been proposed in [40] and further developed in [47]. Here, we ask whether this model outperforms the classical models of tumor growth when fitted with the NSCLC patient data [24].

By fitting the model to the data, we can additionally ask whether different treatment protocols would possibly have been better for some of the patients. The game-theoretic model permits the application of Stackelberg evolutionary game theory by letting the physician anticipate the cancer cell’s evolutionary and ecological dynamics. Besides improving the accuracy for predicting tumor dynamics, modeling cancer therapy as an evolutionary game may help optimizing cancer treatment.

This paper is organized as follows. In Section 2, we introduce the model and fit it with a subset of the anonymized data used in [24]. The subset of patients with metastatic NSCLC were treated with either immune checkpoint inhibition or chemotherapy. In Section 3, we evaluate the results obtained, as well as the accuracy of the model in predicting the future dynamics of the tumor. We compare our results with those given by the classical population growth models explored in [24]. Moreover, we show how our model can be used to design alternative therapies for optimizing any pre-specified treatment goals. We can than use the patient-specific parameterizations to predict outcomes had the patients undergone a different treatment. We can also explore other theoretical scenarios, such as the effect of treatments that target the evolution of resistance. Section 4 concludes by summarizing the main outcomes and discussing limitations and future directions.

## 2 Material and methods

The methods are illustrated in Figure 1 and explained in the following sections.

**Figure 1:**
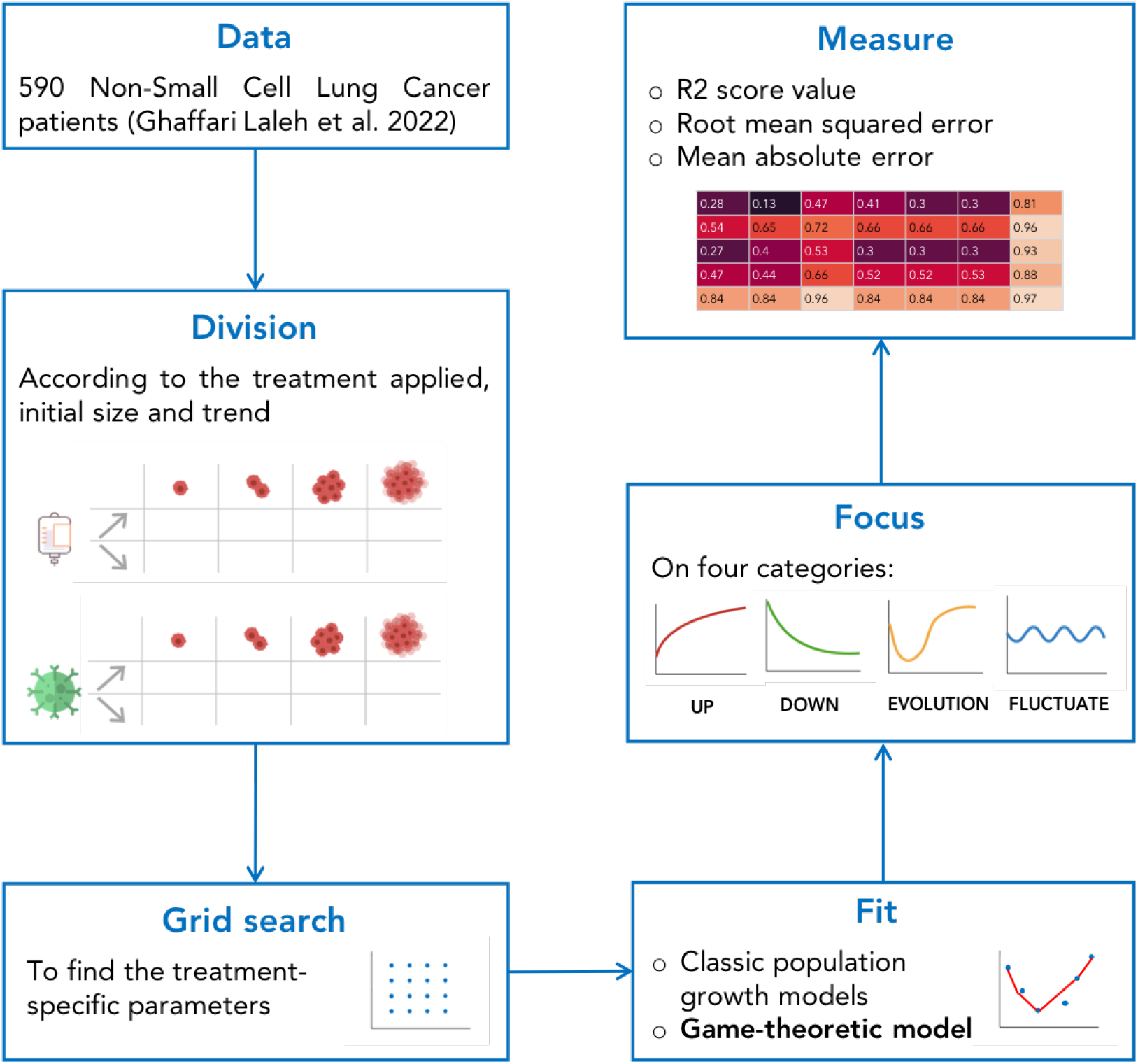
Scheme of the methods. After an initial cleaning and pre-processing of the data, the patients are divided into different clusters based on the initial size of the tumor (Size 1, Size 2, Size 3, Size 4), on the initial behavior (increasing or decreasing) and on the treatment received (chemotherapy or immunotherapy). The treatment specific parameters are selected using a grid search approach. The data are fitted to the classical population growth models analyzed in [24] and to the game-theoretic model. To better highlight the potential of the game-theoretic model we further stratify the patients into four categories (up, down, evolution, fluctuate) and compare the results of the fit in terms of different metrics.

### 2.1 Game-theoretic model

Here we model the cancer cells’ response to treatment as an evolutionary game as in [40]. The physician controls one decision variable *m*, which represents either the dose of Docetaxel, a chemotherapy drug, or an immune checkpoint inhibitor that targets the PD-1 / PD-L1 axis (MPDL3280A). As no patients in our data set received both of the treatments together, we use the same model for both therapeutic options, with a slightly different assumption on the treatment dose *m*. In the case of chemotherapy, the physician can regulate the drug dose and *m* exists on a continuum between 0 and 1, where the standard of care corresponds to *m* = 1 and *m* = 0 means that no treatment is given. On the other hand, in the case of immunotherapy, the physician only has two options: to apply the treatment (*m* = 1) or not (*m* = 0). This is because effect of immunotherapy is not proportional to its dose and is instead either effective or not, as investigated in [36]. For both treatments, when fitting the model to the patient data, we assume that the dose is fixed and *m* = 1 when the treatment is applied and *m* = 0 otherwise. However, when looking for alternative treatments, we consider varying *m* between 0 and 1 in the case of the chemotherapy. We model the cancer’s eco-evolutionary dynamics by means of a population vector *x* of cancer cells, with therapy resistance as a continuous trait *u* ∈ [0, 1], where *u* indicates the intensity of therapy resistance: cells with *u* = 0 are most susceptible (sensitive) to the treatment, while cells with *u* = 1 are maximally resistant to the treatment [45, 47, 40].

In order to integrate the time scales associated with the ecological and the evolutionary dynamics, we use a general approach, called Darwinian dynamics, where the population dynamics and the dynamics of the evolving trait are modeled using a fitness-generating function [10, 53]. We assume that the strategies of the individual cells in the evolving population are inherited and that their payoffs depend on the expected fitness (representing per capita growth rates), which we denote as *G*(*u, x, m*) and refer to as the fitness-generating function, or *G*-function. The ecological dynamics are defined as

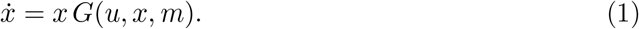

The fitness-generating function also determines the evolutionary dynamics that describe how the cancer cells level of resistance evolves in response to the physician’s treatment choice and are defined as

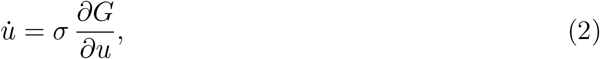

where *σ* determines the evolutionary speed. The G-function approach allows us to determine how natural selection will act on a population’s ecological and evolutionary dynamics. While the function *G*(*u, x, m*) could have different forms, here we confine ourselves to a form used in the existing literature [45, 47, 40]:

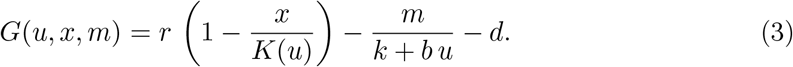

The first term of (3), 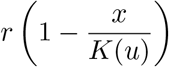, refers to the per capita growth rate of the cancer cells, where *r* represents the intrinsic growth rate, *x* the population of cancer cells, *u* their resistance strategy, and *K*(*u*) their carrying capacity, which is assumed to be a function of the resistance level *u*. In particular, similarly to [60, 11], we assume that there is a cost of resistance manifested in the carrying capacity such that *K*(*u*) decreases as *u* increases: 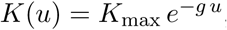, where *K*_max_ is a constant that indicates the maximum possible carrying capacity and *g* determines the magnitude of the cost of resistance. We assume that the maintenance of these resistance mechanisms might require extra energy that cannot be devoted to proliferation, making resistant cells less efficient at utilizing resources for survival and proliferation in the absence of treatment [22]. The second term of the G-function, 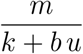, represents the mortality due to the treatment, where *m* is the treatment dose, *k* is innate resistance, and *b* is the benefit of the resistance trait in reducing drug efficacy. The background rate of cell death is given by *d*. Table 1 summarizes all parameters, their definitions and values.

**Table 1:**
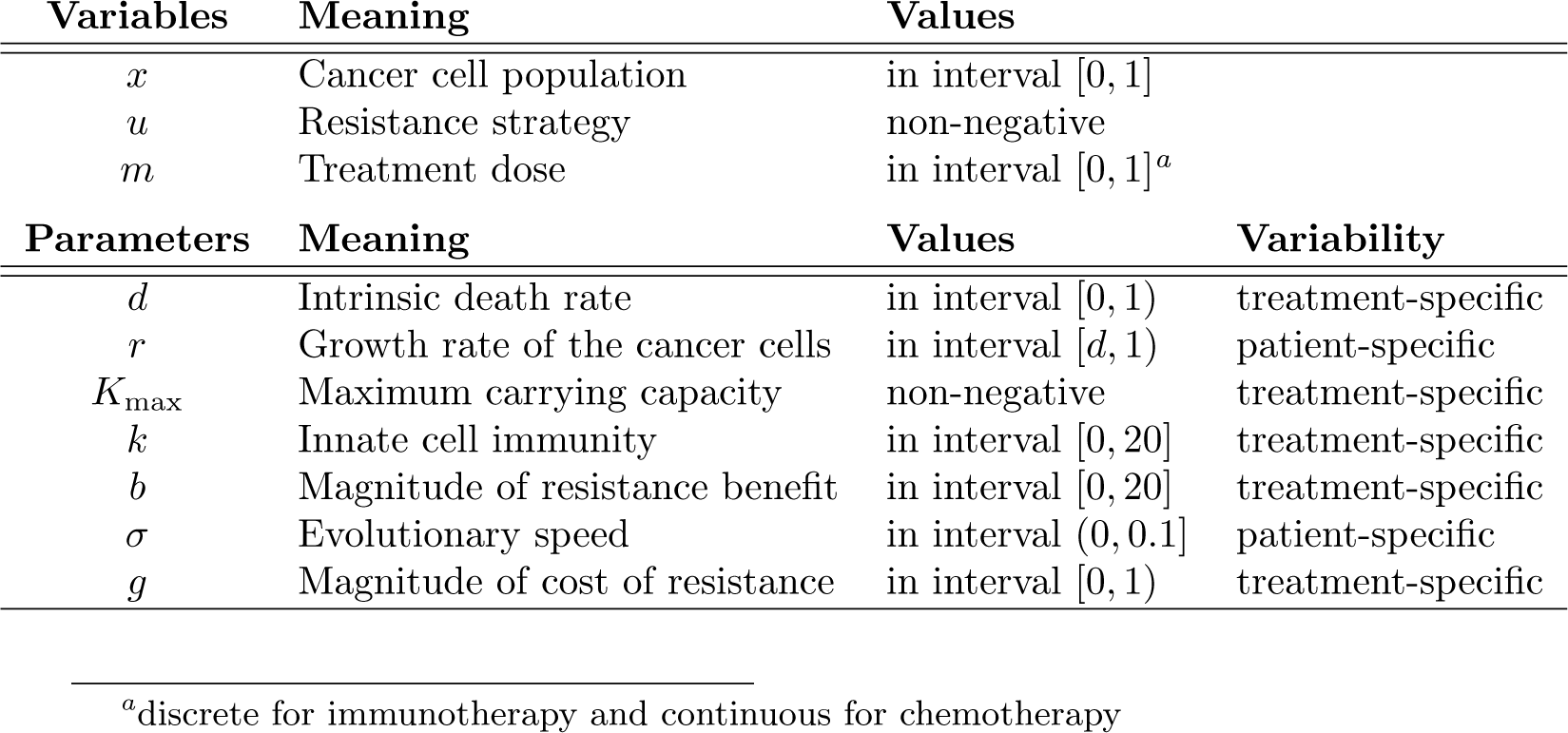
Variables and parameters of the model.

### 2.2 Data and their pre-processing

We used data sets from four different clinical trials reported in [24], which correspond to NSCLC. All the data correspond to patients with metastatic NSCLC treated with Atezolizumab (previously known as MPDL3280A), which is an immune checkpoint inhibitor directed against the Programmed Death Ligand 1 (PD-L1), or with Docetaxel, a chemotherapeutic agent. The data used consists of the one-dimensional longest dimension of the target and non-target lesions manually measured from CT scans. We cleaned and pre-processed the data following the guidelines in [24] in order to make comparable results.

We selected the primary tumor (target lesion) of each patient and considered only those tumors for which at least 6 temporal observations are available. This enables a more robust fitting of the mathematical model to the time series data. This process resulted in a cohort of 590 patients (Figure 1). Patients varied in the number of measurement time points for the target lesion. The distribution of patients among studies and treatment arms is explained in more detail in [24].

In order to estimate the population of cancer cells of each tumor from the measurements of the longest diameter (*LD*), the tumor volume (*V*) was calculated using the formula *V* = 0.5 *·* (*LD*)^3^, following common practice in tumor modelling when only a single tumor measurement is available [20].

Furthermore, as the range of tumor volumes was very large, between 0 and 5926176 mm3, the values were rescaled and normalized to range from 0 to 1, using the formula

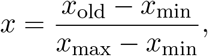

where *x* represents the rescaled volume, *x*_old_ is the original volume estimated from the diameter, and *x*_min_ and *x*_max_ represent the minimum and maximum volumes from the patient’s time series, respectively. This was done in order to have only three patient-specific parameters in the model, while fixing other parameters to the same value for all of the patients.

To present the results more effectively, the tumors were categorized into four treatment trend categories: “Up”, “Down”, “Fluctuate” and “Evolution” (Figure 1). To do this, for each tumor we created a vector containing the differences between each *LD* measurement at time point *t*_*k*+1_ and the previous measurement at time point *t*_*k*_, for all the time points where a measurement was taken. If the *LD* at *t*_*k*+1_ is bigger than at *t*_*k*_, we consider the difference as positive and vice versa. The tumors will be classified and categorized as follows:

- The “Up” category includes patients whose difference vector values are always positive and patients with a positive difference after the first measurement if the ratio between the sum of all positive values to the sum of all negative values is greater than 2.
- The “Down” category includes patients whose difference vector values are always negative or a negative difference after the first measurement if the overall ratio between the sum of all negative values to the sum of all positive values is greater than 2.
- The “Evolution” category includes patients who initially respond to treatment and then start experiencing tumor progression. Here the evolution of resistance is most pronounced. We include tumors for which the first two values of the difference vector are negative, the maximum of the volume corresponds to the first or the last time point, and the sum of the last two values of the difference vector is greater than −1 times the first value of the vector divided by 2 (i.e., the evolution of resistance causes the tumor to grow at least half as much as it shrank at the start of the treatment).
- The “Fluctuate” category contains all patients who correspond to none of the previous categories.

The categories “Up” and “Down” correspond to those described in [24], while the category “Fluctuate” defined in [24] is now divided into two categories: “Evolution” and “Fluctuate”. This is done because tumors in the category “Evolution” are more likely to exhibit a U-shape characteristic of the evolution of treatment-induced resistance. Since our model takes evolution of therapy resistance explicitly into account, we want to confirm the hypothesis that it fits the tumors in this category more accurately than population growth models absent evolution.

### 2.3 Fitting the model

The model serves to estimate the parameters introduced in Table 1 by fitting to the values of *x* at the different time points.

Parameter estimation was done using GEKKO v0.2.8, a Python package for machine learning and optimization, specialized in dynamic optimization of differential algebraic equations [4]. As there are seven free parameters (*d, r, K*_max_, *k, b, σ* and *g*), estimating their values poses a significant challenge. We let two of the parameters and the initial condition for *u* to be patient-specific. We fixed the values of the remaining parameters. Initial experimentation suggested that letting *r* and *σ* be patient-specific and fixing the remaining parameters provided some of the best fits.

Thus, to fit the model, the parameters *b, k, K*_max_, *g*, and *d* were fitted with the assumption that they are treatment-specific, but otherwise the same for all patients receiving the same treatment. We then let *σ, r* and the initial value of *u* be patient-specific, as summarized in Table 1. They are estimated separately for each tumor with the objective of minimizing the error between the real data and the predictions given by the model. For all tests, we assume *m* = 1 when treatment is applied.

The values of the parameters which are not patient-specific are selected after performing a grid search to find the combination that gives the best results. The model was fitted to the data for each tumor for each combination of parameters and then, the mean squared error was evaluated per tumor. The final value of the average error of all the tumors was then used to select the best parameters.

### 2.4 Separating tumors into different clusters

Due to the fact that the behaviours and volumes of the tumors vary significantly from patient to patient and the number of parameters to optimize is relatively large, we separate the tumors into different clusters such that each cluster has different parameter values.

First of all, as we mentioned before, the parameters *b, k, K*_max_, *g*, and *d* are assumed to be treatment-specific, so different values are estimated for each of the two treatments. Within each treatment group, we further stratify the patients according to the following criteria. The first clustering criterion is the initial volume of the tumors, as the range of the volumes is large and the volume of the tumor might have an impact on its evolution, and thus on the parameters. Moreover, this information can be extracted solely from the first data point, so serves immediately when predicting future tumor sizes.

The second clustering criterion is the initial trend of the tumors, that is, whether they increase or decrease once treatment starts. This is a reasonable criterion as patients are clinically categorized as non-responders based on an initial increase, and as responders based on an initial decrease. Furthermore, the first reaction to treatment may contain valuable information concerning future tumor dynamics as measured from these first two data points. Therefore, for each treatment group, tumors are separated into four clusters according to initial volume (rescaled and normalized), Size 1, Size 2, Size 3 and Size 4, and two sub-clusters according to the initial behavior, increasing and decreasing, so in total we differentiate eight groups per treatment, corresponding to all the possible combinations of initial trend and initial volume (Figure 1). Tumors were classified as Size 1 if their volume was smaller than 1021.9 mm^3^, Size 2 if their volume was between 1021.9 mm^3^ and 10219 mm^3^, Size 3 if it was between 10219 mm^3^ and 61314.24 mm^3^ and Size 4 if the volume surpassed the latter threshold. Tumors were included in the increasing group if the second measurement of the diameter was larger than or equal to the first one, and in the decreasing group if it was smaller.

For each group, a grid search was performed to fit the parameters, where for each grid point patient-specific fit was carried out (Figure 1). The parameters that lead to the best fit per group were selected. Their values are summarized in Table 2.

**Table 2:**
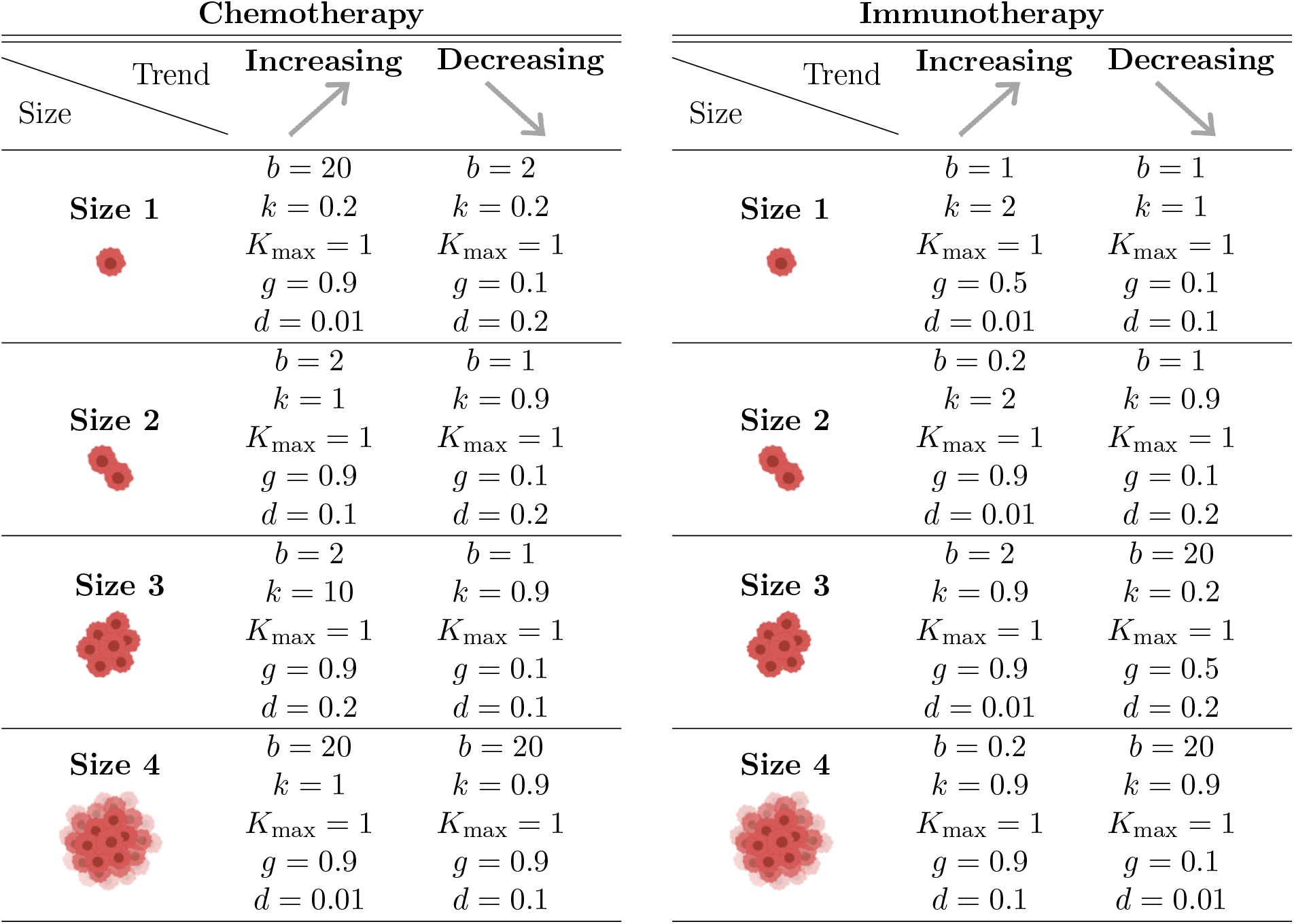
The values of treatment-specific parameters fitted using grid search. We cluster the patients according to the initial tumor volume (Size 1 ≤ 10219 mm^3^, 10219 mm^3^ < Size 2 ≤ 10219 mm^3^, 10219 mm^3^ < Size 3 ≤ 61314.24 mm^3^ and Size 4 *>* 61314.24 mm^3^), the initial behavior (increasing or decreasing) and the treatment received (chemotherapy or immunotherapy).

| Chemotherapy |  |  |  |
| --- | --- | --- | --- |
| Trend |  | Increasing | Decreasing |
| Size |  |  |  |
| Size 1 | | $b = 20$ | $b = 2$ |
| | | $k = 0.2$ | $k = 0.2$ |
| | | $K_{\max} = 1$ | $K_{\max} = 1$ |
| | | $g = 0.9$ | $g = 0.1$ |
| | | $d = 0.01$ | $d = 0.2$ |
| Size 2 | | $b = 2$ | $b = 1$ |
| | | $k = 1$ | $k = 0.9$ |
| | | $K_{\max} = 1$ | $K_{\max} = 1$ |
| | | $g = 0.9$ | $g = 0.1$ |
| | | $d = 0.1$ | $d = 0.2$ |
| Size 3 | | $b = 2$ | $b = 1$ |
| | | $k = 10$ | $k = 0.9$ |
| | | $K_{\max} = 1$ | $K_{\max} = 1$ |
| | | $g = 0.9$ | $g = 0.1$ |
| | | $d = 0.2$ | $d = 0.1$ |
| Size 4 | | $b = 20$ | $b = 20$ |
| | | $k = 1$ | $k = 0.9$ |
| | | $K_{\max} = 1$ | $K_{\max} = 1$ |
| | | $g = 0.9$ | $g = 0.9$ |
| | | $d = 0.01$ | $d = 0.1$ |

Table 2: The values of treatment-specific parameters fitted using grid search.
| Immunotherapy |  |  |  |
| --- | --- | --- | --- |
| Trend |  | Increasing | Decreasing |
| Size |  |  |  |
| Size 1 | | $b = 1$ | $b = 1$ |
| | | $k = 2$ | $k = 1$ |
| | | $K_{\max} = 1$ | $K_{\max} = 1$ |
| | | $g = 0.5$ | $g = 0.1$ |
| | | $d = 0.01$ | $d = 0.1$ |
| Size 2 | | $b = 0.2$ | $b = 1$ |
| | | $k = 2$ | $k = 0.9$ |
| | | $K_{\max} = 1$ | $K_{\max} = 1$ |
| | | $g = 0.9$ | $g = 0.1$ |
| | | $d = 0.01$ | $d = 0.2$ |
| Size 3 | | $b = 2$ | $b = 20$ |
| | | $k = 0.9$ | $k = 0.2$ |
| | | $K_{\max} = 1$ | $K_{\max} = 1$ |
| | | $g = 0.9$ | $g = 0.5$ |
| | | $d = 0.01$ | $d = 0.2$ |
| Size 4 | | $b = 0.2$ | $b = 20$ |
| | | $k = 0.9$ | $k = 0.9$ |
| | | $K_{\max} = 1$ | $K_{\max} = 1$ |
| | | $g = 0.9$ | $g = 0.1$ |
| | | $d = 0.1$ | $d = 0.01$ |

### 2.5 Experiments

#### Comparing the fit of the game-theoretic model to that of the classical models

In experiments, we evaluated how well the game-theoretic model defined by equations (1)–(2) matched the tumor volume trajectories of patients subject to immunotherapy or chemotherapy and compared the results to the ones obtained with the classical population models explored in [24] (Figure 1). In order to compare the performance between the game-theoretic model and the six population models explored in [24], we calculated different metrics of the goodness of fit (for each of the study arms and for each of the trend categories), namely the R2-score, the root mean squared error (RMSE) and the mean absolute error (Figure 1). Further, we compare the fit of the model with and without evolution of resistance to see whether a possible better fit was simply due to more parameters in the game-theoretic model, or rather, due to its allowing for therapy resistance to evolve.

#### Predicting treatment response based on initial data points only

After fitting values of the treatment-specific parameters of each group, we test whether the model can predict the later data points from the early treatment response. More specifically, if we have *n* data points, we use the first *n* − 3 points to estimate the values of the patient-specific parameters, *r, u*(0) and *σ*, and then we solve equations (1) and (2) to get the value of *x* at the time of the last measurement. Then, we evaluate the result by computing the mean absolute error between the predicted volume at time *t*_*n*_ and the real value.

#### Optimizing a pre-defined treatment objective

Here we expand our model into a Stackelberg evolutionary game [46, 58, 50, 47] where the physician as a rational leader tries to optimize a predefined objective, by changing the timing of giving or not-giving the drug, and, in the case of chemotherapy, also the dose level. Different objective functions can be explored, such as minimizing the final tumor burden, minimizing the tumor burden at every time point, minimizing the variance of the volume of the tumor, etc.

We consider the objective of minimizing the final tumor burden, which corresponds to solving the following optimization problem:

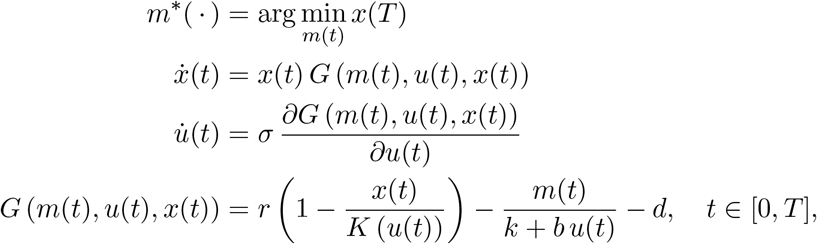

where *T* is the final point considered. For patients subject to immunotherapy, we assume that we can decide whether treatment is on or off following every measurement time point, and so, at each of the time points, the optimal value of *m* ∈ {0, 1} is computed. On the other hand, for patients that received chemotherapy, we consider *m* ∈ [0, 1] and compute the optimal dose following each of the measurement time points.

#### Simulating a different treatment scenario

Since we have assumed that there is a set of parameters that are treatment-specific, namely *K*_max_, *b, d, g* and *k*, we can use them to simulate what would have happened if the patients who received immunotherapy were being given chemotherapy and vice versa.

To do so, we assume that the initial value of the resistance to treatment is the same for both treatments. We take the parameters *K*_max_, *b, d, g* and *k* estimated for a certain treatment and the parameters *u*(0), *r*_max_ and *σ* estimated for a certain patient, and we simulate the ecological and evolutionary dynamics of the tumor subject to this treatment.

Thus, for patients who received Docetaxel, we simulated what would have happened if they had received immunotherapy, and vice versa.

#### Simulating a treatment targeting the evolution of resistance

Here we assume there exists a treatment that targets evolution of resistance, that is, that makes *u* decrease instead of increase, and explore how application of such a drug would affect tumor evolution and tumor dynamics.

## 3 Results

### 3.1 Comparing the fit of the game-theoretic model with classical ODE models

As shown in Figure 2, Figure 3 and Figure 7, in general the game-theoretic model provides a better fit for both metrics for most of the groups, although the difference in performance varies with category. Indeed, for the categories “Up”, “Down” and “Fluctuate”, the R2-score values are often similar to the best scores obtained from the other models. For tumors in the “Evolution” category, the game-theoretic model significantly outperforms the other models in most of the treatment arms. In fact, with the game-theoretic model we obtain an R2-score higher than 0.8 for all arms except one, compared to values below 0.6 with the classic ODE population models in most cases. This is confirmed by the other error measures considered. This indicates that the game-theoretic model is indeed more suitable for describing tumors that exhibit an evolution of resistance to treatment, featuring a U-shape in their trend of growth over time. Figure 4 shows two fits obtained for tumors in the category “Evolution”, that could not be suitably fitted with the classical ODE models.

**Figure 2:**
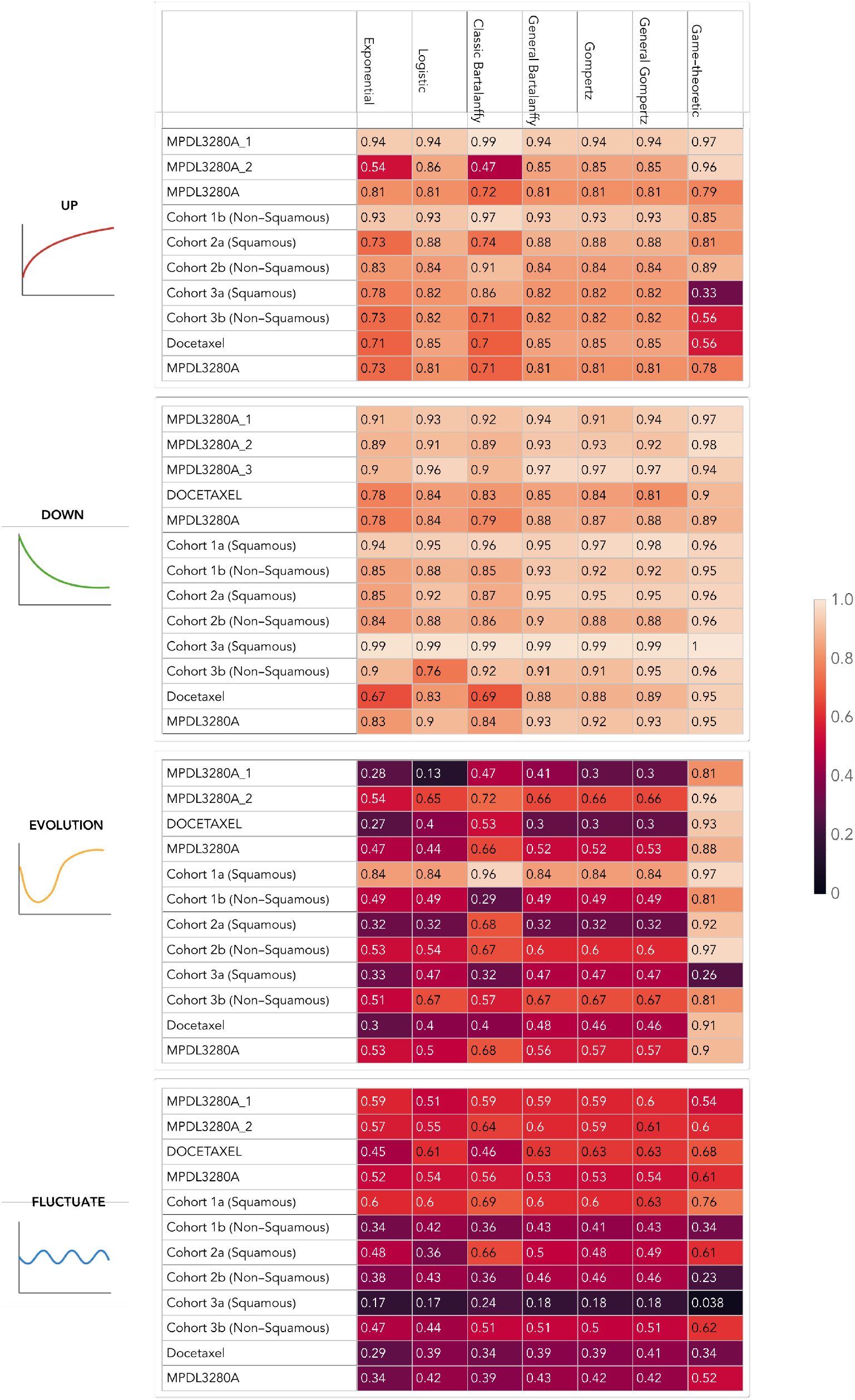
R2 score values for each model and each trend category, where each model was fitted separately for that category. A higher value corresponds to a better fit.

**Figure 3:**
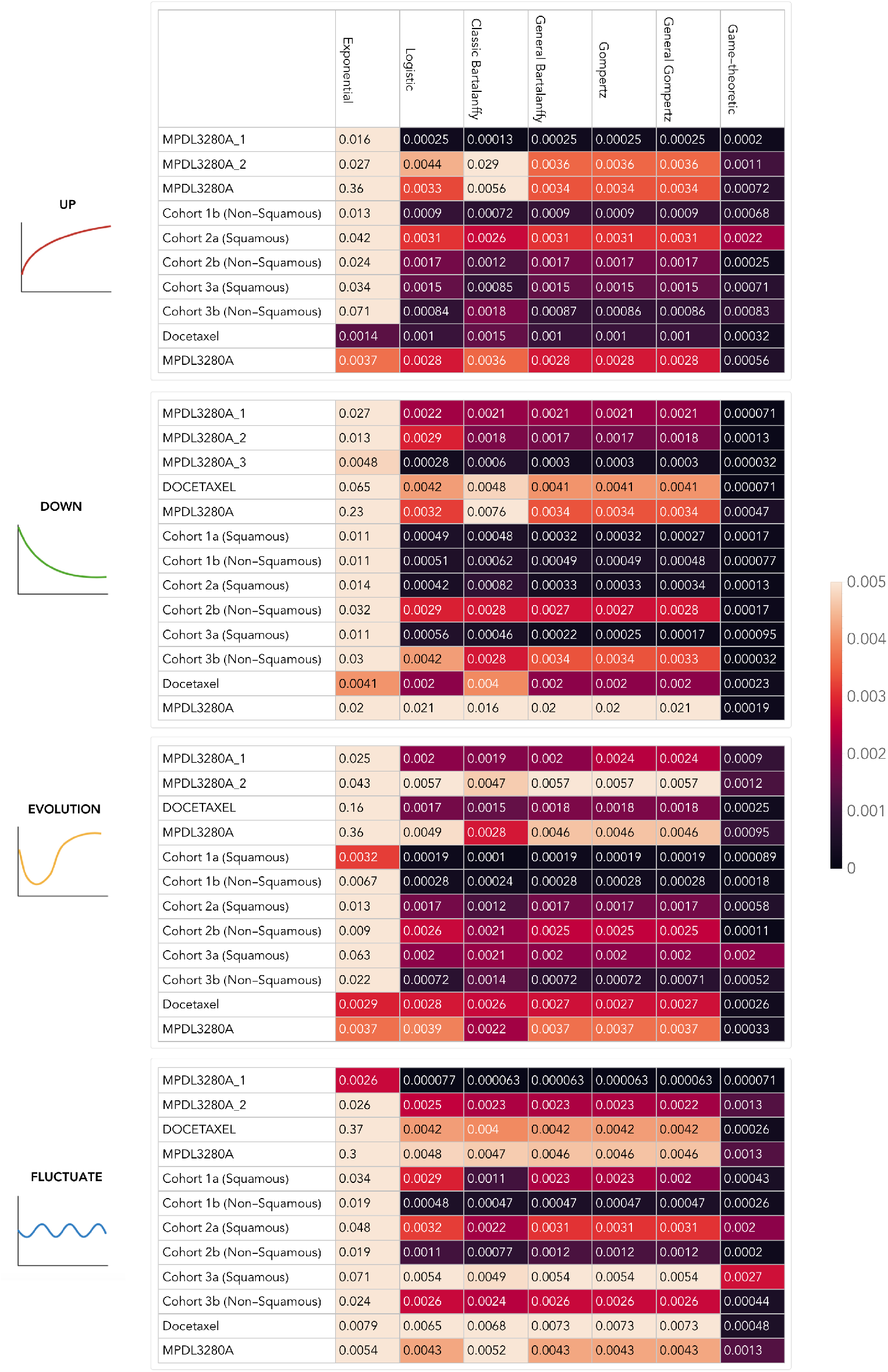
Root mean squared error for each model and each trend category, where each model was fitted separately for that category. A lower value corresponds to a better fit.

**Figure 4:**
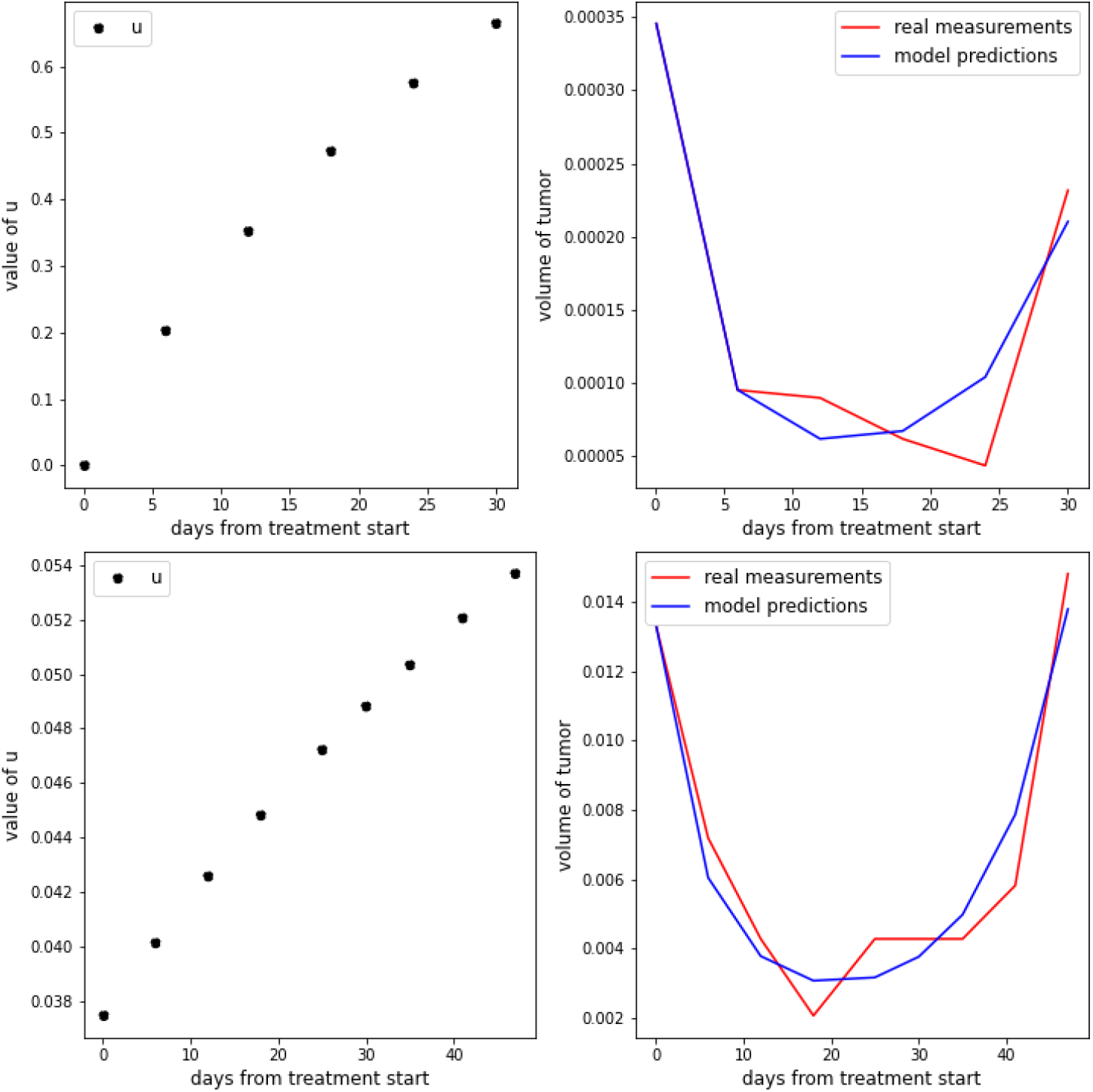
Two examples of the fits obtained for tumors in the “Evolution” category for a patient treated with Docetaxel (top) and a patient treated with immunotherapy (bottom).

On the other hand, the R2-scores for tumors in the trend category “Fluctuate” are below 0.6 in most of the groups for all of the models, which indicates a systematic limitation to accurately fit tumors that experience pseudoprogression, i.e., tumors that keep growing for a relatively long period of time after the start of treatment and then start responding to treatment, or tumors that grow and decrease alternatively over time.

According to our results, the fits given by the game-theoretic model are generally better than those of the classic population models. In order to determine whether this superiority is purely due to the higher complexity of the model or also influenced by the assumption of including resistance as an evolving trait, we compared the results obtained with the game-theoretic model when we assume there is no evolution of resistance (*σ* = 0) with those obtained when we assume that evolution of resistance occurs (*σ* > 0).

Figure 5 shows that, generally, tumors that exhibit a monotonic behaviour, are fitted well under the assumption that resistance does not evolve. However, for tumors that exhibit changing trends over time – typically presented an initial response to treatment followed by an increase of volume over time – we obtain a poor fit when *σ* = 0 (Figure 5). These cases support the importance of modeling resistance as an evolving trait.

**Figure 5:**
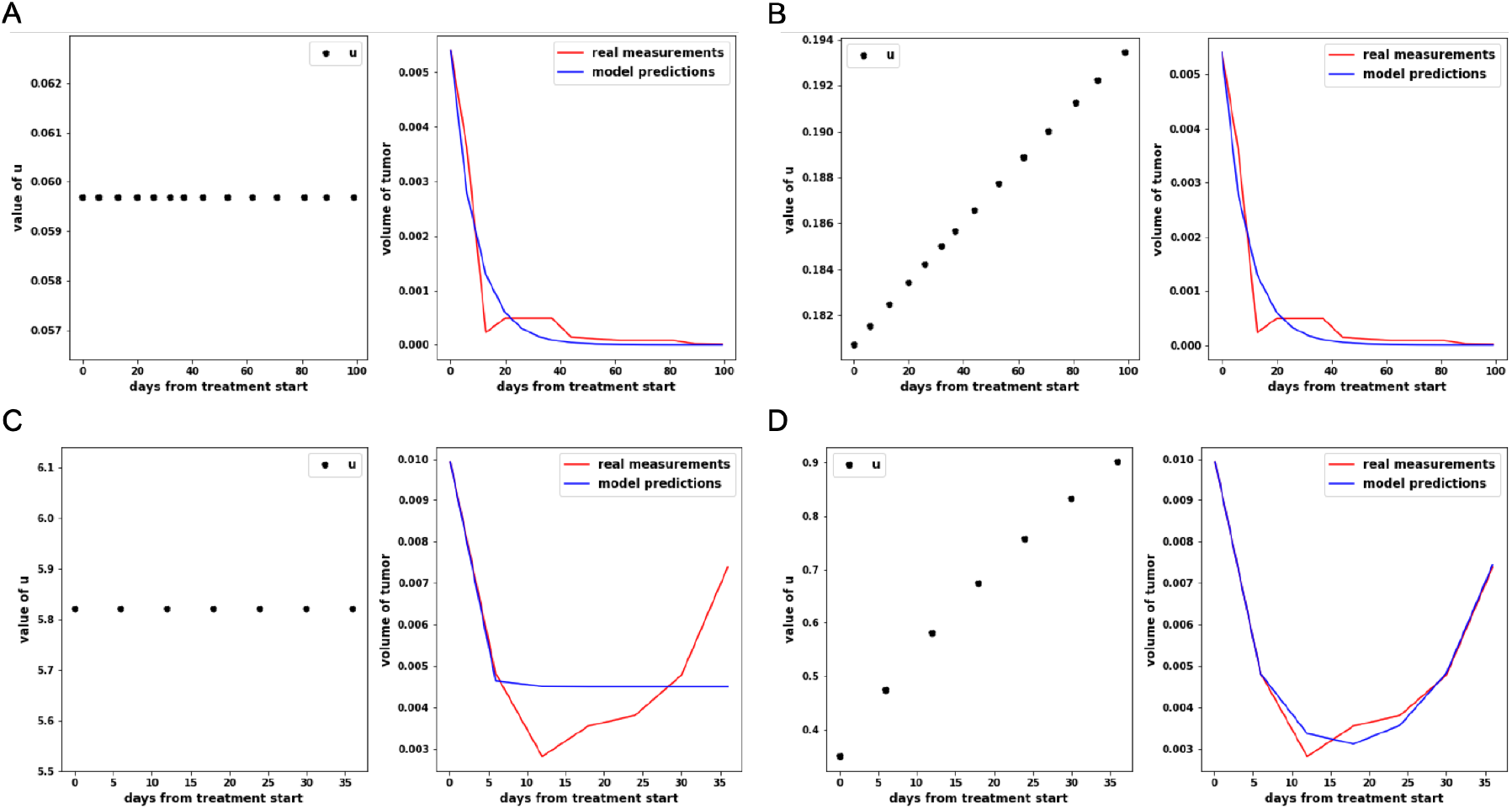
Fits obtained assuming: (A) no evolution of resistance for a patient treated with immunotherapy; (B) evolution of resistance for a patient treated with immunotherapy; (C) no evolution of resistance for a patient treated with Docetaxel; (D) evolution of resistance for a patient treated with Docetaxel.

### 3.2 Predicting treatment response based on initial data points

When comparing how the models are able to predict the later data points from the early treatment response, we noticed that, in general, we obtain a lower mean absolute error with the game-theoretic model than with the ones used in [24] (Figure 7). Of particular interest is the case of U-shaped tumors, that could not be well predicted by any of the classic ODE models explored in [24]. For most of these tumors, the game-theoretic model provides a more accurate prediction (Figure 6 shows two examples), although in some cases the estimated growth due to evolution of resistance is considerably higher than the real growth, while in some other cases, where resistance appears at a later time, the model fails to predict this growth.

**Figure 6:**
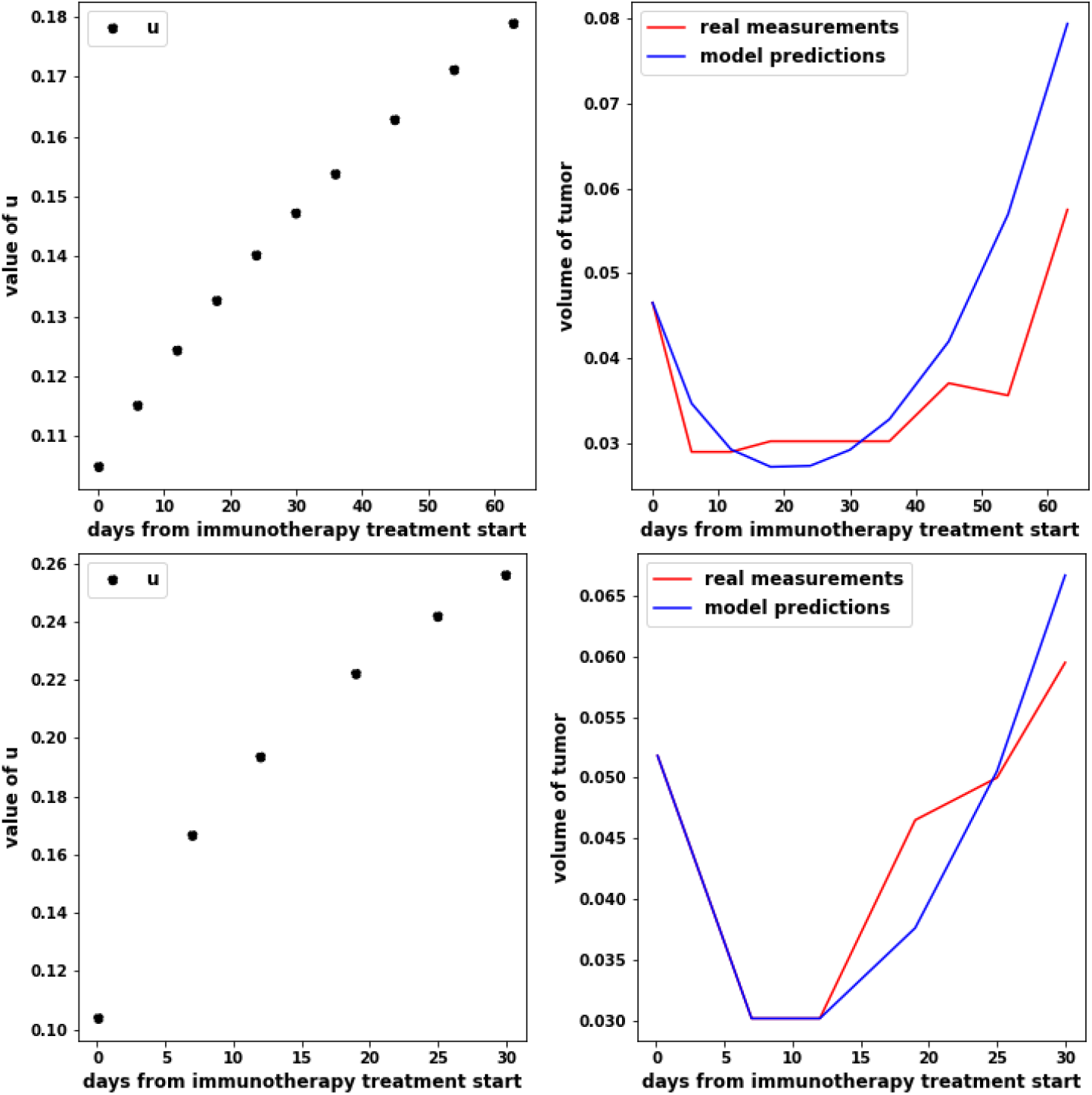
Predictions given for two tumors in the “Evolution” category for a patient treated with immunotherapy (top) and a patient treated with Docetaxel (bottom).

### 3.3 Optimizing a pre-defined treatment objective

Simulations show that in general it is not possible to stabilize the tumor volume, and that the optimal strategy is usually to treat patients continuously at maximum tolerable dose. This is partially due to our objective being the final tumor burden. For a limited number of patients we find that a different treatment schedule would lead to the same final tumor burden. For example, Figure 8 shows the case of two patients who received Docetaxel. Lowering the dose of treatment yields the same final tumor burden as applying constant maximum tolerable dose, although the tumor volume at previous time points is higher. If the tumor burden outcome is the same, then a lower treatment dose will correspond to a higher quality of life given drug toxicities and discomforts caused by the medication [1, 55]. Furthermore, if we simulate the evolution of the tumor for a longer period of time, aiming at minimizing the tumor burden at the final time point, we find that for a limited number of cases allowing the tumor to grow at the beginning results in a lower final tumor burden (Figure 9). Nevertheless, when following this treatment schedule, the tumor tends to grow until it has reached a volume close to the maximum over all of the patients from the dataset. The challenge in designing an evolutionary treatment for these patients is that the growth rate of stage 4 NSCLC tumors is too high to allow for periods of unrestricted tumor growth [16, 15].

**Figure 7:**
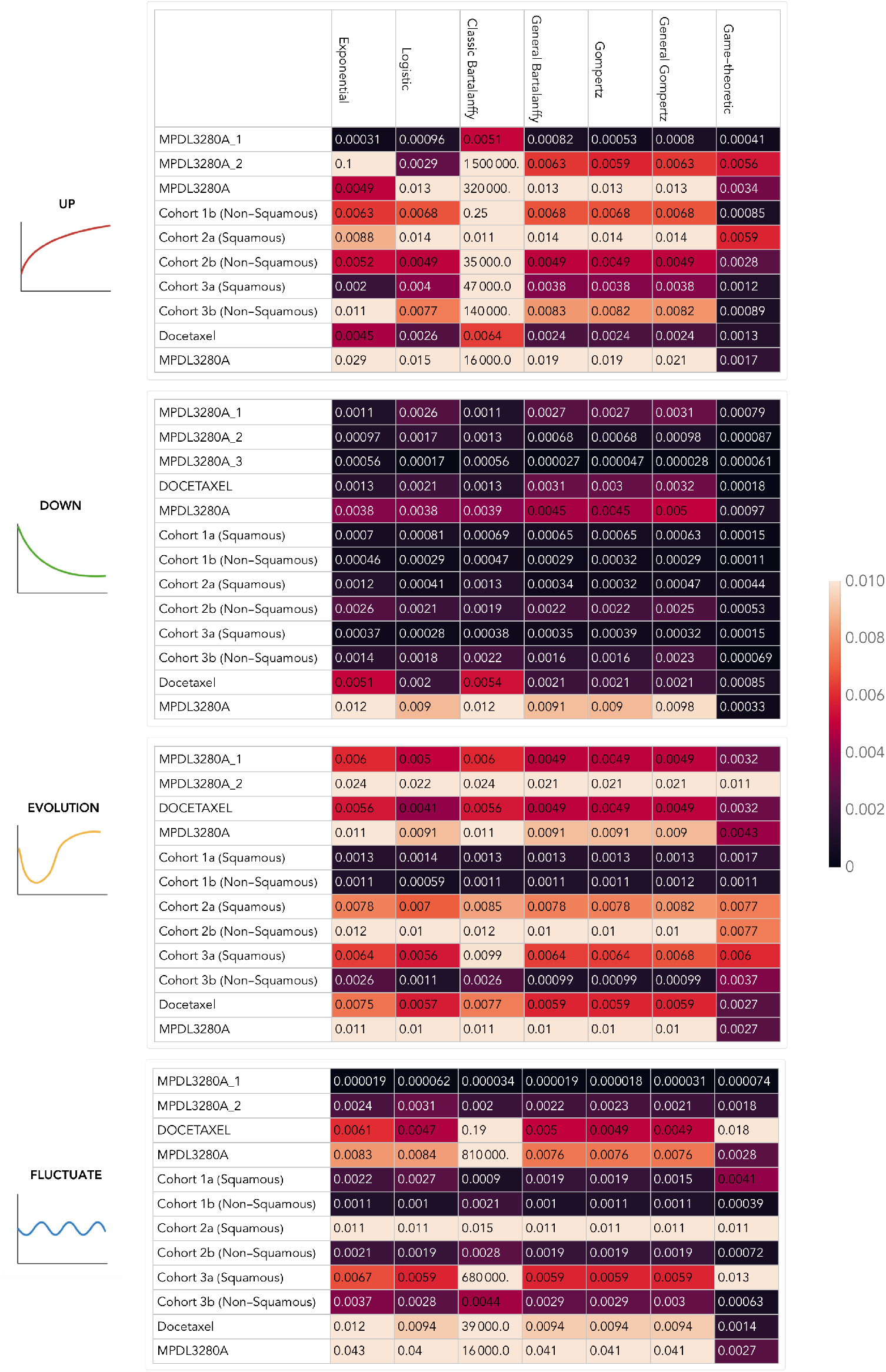
Mean absolute error values for prediction of tumor dynamics, fitted separately per model and trend category. A lower value corresponds to a better fit.

**Figure 8:**
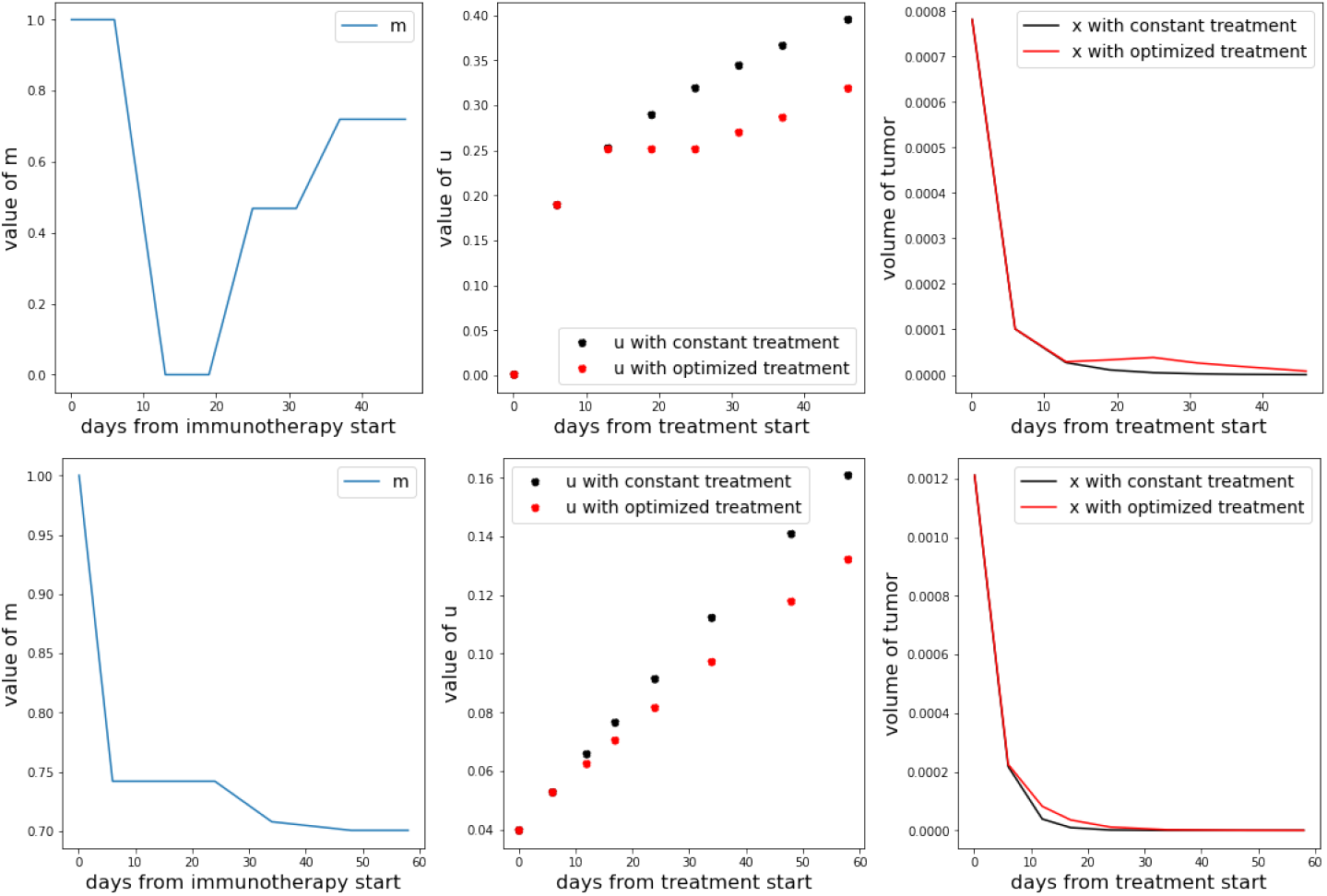
Comparison between continuous treatment and optimized treatment (with the aim of minimizing the final tumor burden) in a patient treated with Docetaxel.

**Figure 9:**
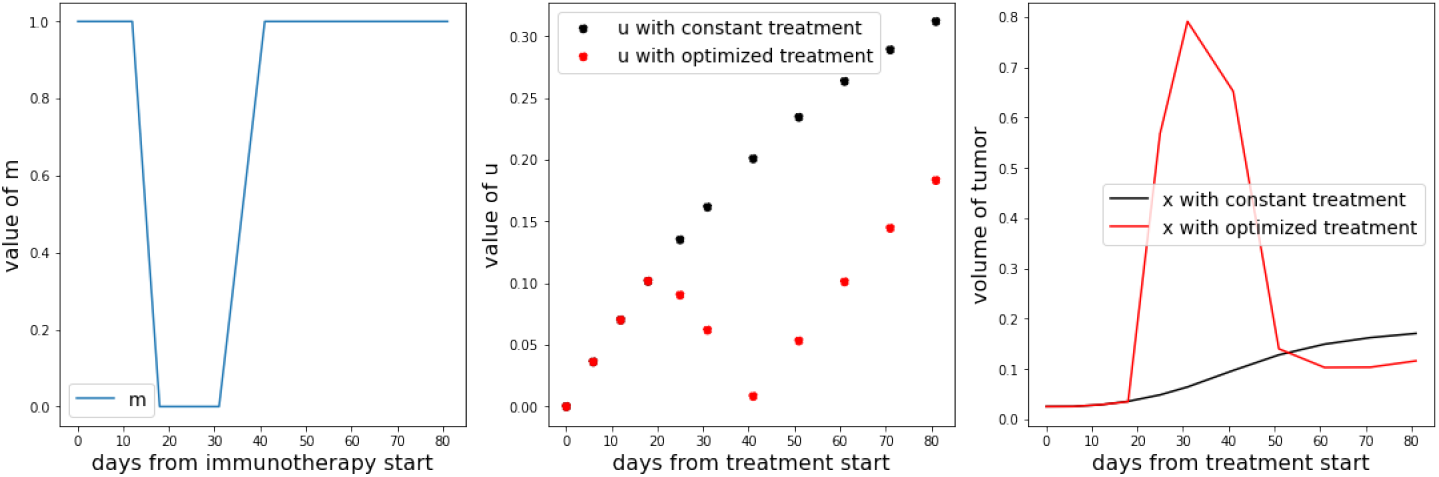
Comparison between continuous treatment and optimized treatment on a longer time period (with the aim of minimizing the final tumor burden) in a patient treated with immunotherapy.

On the other hand, the treatment objective here targets the tumor burden only, thus it may be that optimizing an overall quality of life metric will lead to superior results from an evolutionary therapy that successfully anticipates and steers therapy resistance.

### 3.4 Simulating a different treatment scenario

For patients who received Docetaxel, we simulated what would have happened if they had received immunotherapy. We found that the majority remains better off as 75% of these patients obtained a lower final tumor burden with Docetaxel than with immunotherapy. Figure 10 shows the comparison between the real tumor evolution of a patient subject to Docetaxel and the simulation of the eco-evolutionary dynamics under immunotherapy, which would lead to a noticeably higher final tumor burden.

**Figure 10:**
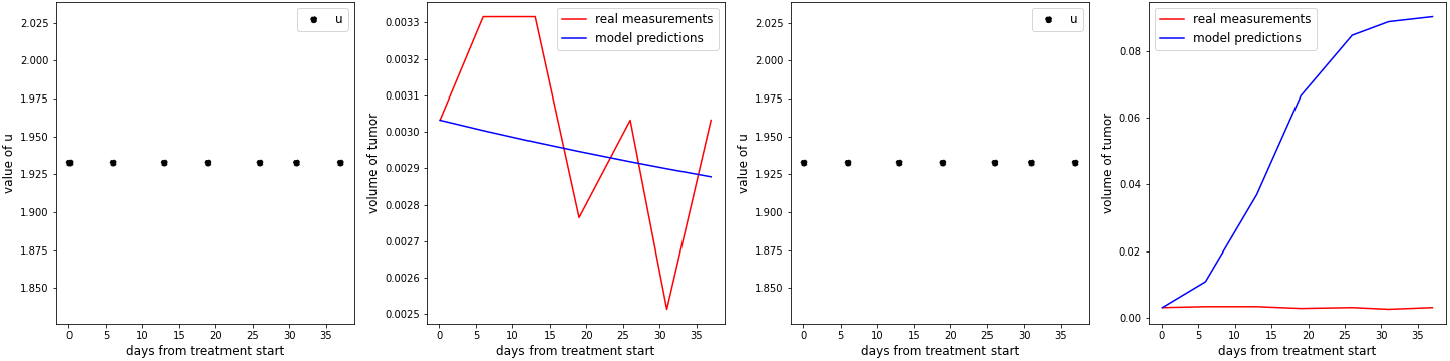
Real tumor evolution subject to Docetaxel (left) versus simulation of immunotherapy (right).

On the other hand, when we simulated the behaviour under Docetaxel for patients who were treated with immunotherapy, the percentage of patients that obtained a lower final tumor burden with their given treatment dropped to 50%. Figure 11 shows an example of a tumor that would have responded better to Docetaxel than to immunotherapy according to simulations.

**Figure 11:**
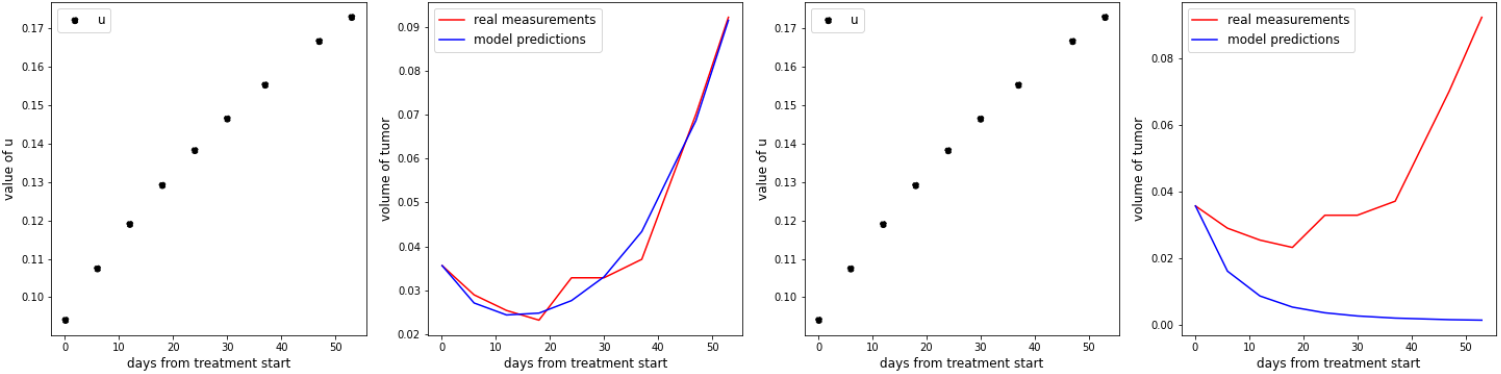
Real tumor evolution subject to immunotherapy (left) versus simulation of Docetaxel (right).

### 3.5 Impact of a treatment targeting the evolution of resistance

If we assume the existence of a treatment that targets evolution of resistance, we find that 58% of the patients would achieve a lower tumor burden than with their actual treatment (Figure 12 reports two such examples). Moreover, the percentage of tumors such that the final tumor burden is lower than the starting one is 75%, compared to 63% with the existing treatments. This alternative treatment is especially useful for tumors in the trend categories “Up” and “Evolution”. More than 70% of the tumors in these categories respond better to the alternative treatment than to traditional treatments, meaning that the final tumor burden was smaller.

**Figure 12:**
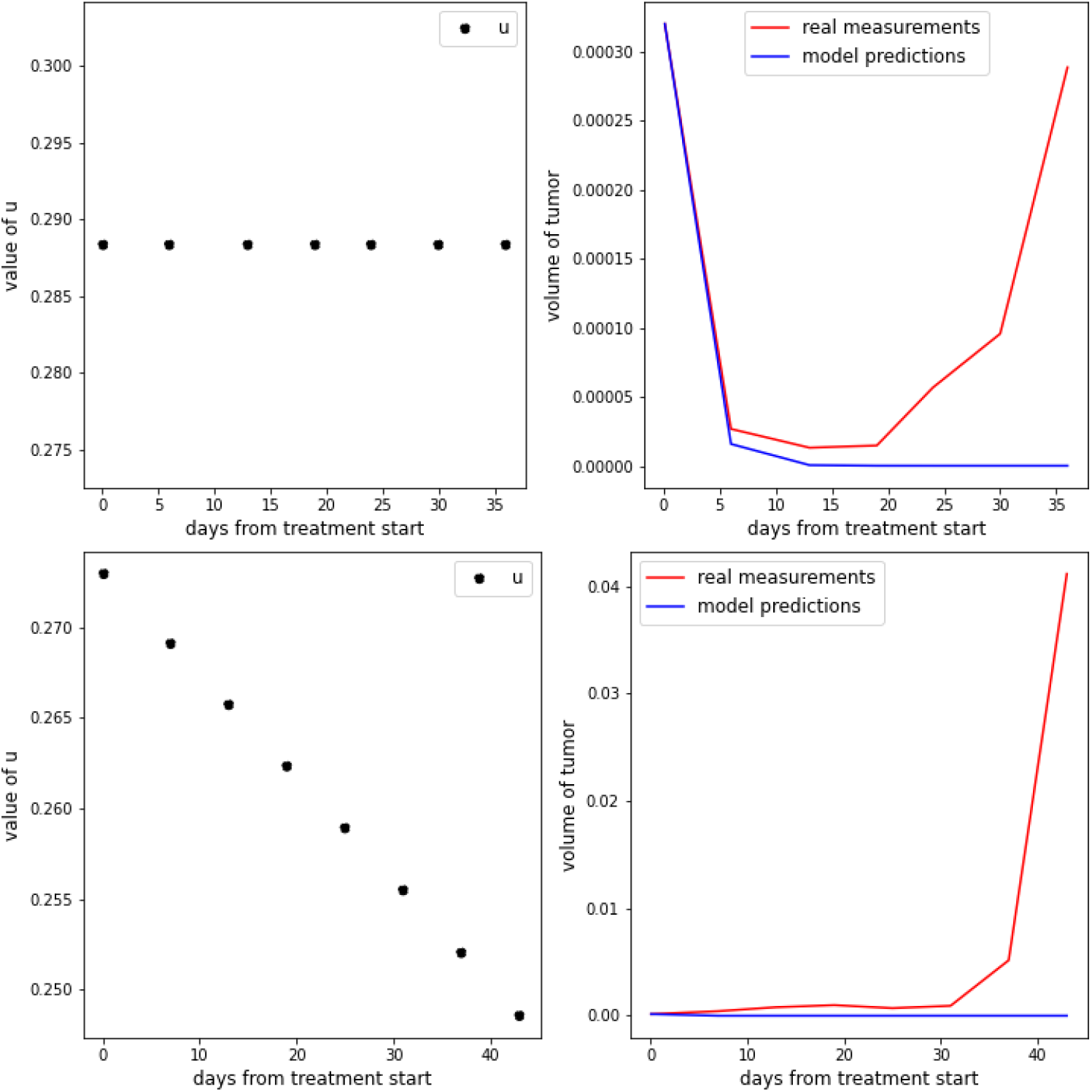
Evolution of tumor volume of two tumors in the category trend “Evolution” under a hypothetical alternative treatment that targets evolution of resistance.

## 4 Discussion

The game-theoretic model introduced here specifically considers the evolution of resistance by the cancer cells in response to therapy. By doing so, it generally provides a good fit to a cohort of patients with metastatic Non-Small Cell Lung Cancer being treated with either an immunotherapy or a chemotherapy. In particular, the model provides a better fit than that which had been achieved by earlier study using standard models of population growth that do not include an evolutionary component [24]. The more accurate fit by the game-theoretic model was most striking relative to the classic population ODE models when tumors exhibited a U-shape in their temporal dynamics. This is not surprising since the population growth models always provide a monotonic trend of tumor burden dynamics under therapy. But, it is important for two reasons. First, such U-shaped dynamics are typical of cancers that first respond and then progress due to the evolution of resistance. Second, the game-theoretic model was able to capture this trajectory as well as provide an estimate of the level of resistance throughout the course of treatment.

Similar models have been proposed for modelling resistance as a continuous trait to fit patient data. The Tumor Growth and Inhibition (TGI) model [9, 25] can have as few as three parameters including tumor cell growth rate, initial drug efficacy at reducing the growth rate, and then a parameter describing how drug efficacy declines with time during therapy. A variant of this is in use by Moffitt’s Evolutionary Tumor Board [43]: the Growth, tumor Death, evolution of drug Resistance, and drug re-Sensitization (GDRS) model. The GDRS, like our model, considers the evolution of increased drug sensitivity when therapy is removed. The TGI model does not. In these three models the difference lies in the formulation of how changing the resistance strategy alters drug efficacy. All three can generate a U-shaped dynamic for tumor growth, and the declining phase and increasing phase of the dynamics can be asymmetric. This separates these models from a simply second order polynomial fit with an upward parabola. Finally, unlike the TGI and GDRS models, our model explicitly considers resistance as a trait that is than linked to its consequences for drug efficacy.

A second class of models sees resistance as evolving from distinct, qualitatively different, cancer cell populations, generally binned as sensitive versus resistant cells [3, 12, 56, 33, 26, 52, 54, 34, 6, 51, 60]. In our model this is akin to introducing two to several cancer subpopulations with fixed, non-evolving, values for their resistance traits. Pressley et al. (2021) provide a hybrid version where one subpopulation’s trait is fixed while the other evolves as a continuous trait [40]. Such models can also predict a U-shaped trajectory of tumor dynamics under therapy, but generally requiring more parameters, and they have been used extensively to model adaptive therapy including applications to patients with prostate cancer and melanoma. An interesting feature emerging from models that view resistance as a qualitative versus quantitative trait concerns the efficacy of adaptive therapy. Generally it seems more effective when the former than latter. And indeed, Pressley et al. found that adaptive therapy regimens were generally no better and sometime inferior to continuous dose therapy.

It remains an empirical question when resistance traits are qualitative versus quantitative. Examples of each exist. In prostate cancer, the cell types are qualitatively different such as cells that require testosterone, those independent of testosterone or those producing their own testosterone [31, 37]. In breast cancer, a form of resistance to hormonal therapy can be quantitatively increasing the production of estrogen receptors [27], or qualitatively rewiring metabolic pathways to be completely independent of estrogen [59]. Efflux pumps, a quantitative trait, confers resistance by binding to and removing chemical agents from the cancer cell, thus conferring multi-drug resistance to such drugs as docetaxel, paclitaxel, methotrexate and others in a variety of cancers [39, 21, 17]. Our model can serve for both quantitative and qualitative resistance mechanisms.

The quality of our model’s fit to the patient data varied both between and within patient categories. Several factors can improve fit. First, including the initial response into the model (using two data points to initialize the efficacy parameter) yielded a lower error, although the results obtained show that in some cases predictions are still not satisfactory. More accurate predictions could be achieved by reducing the number of patient-specific parameters or by providing more information about their values, i.e., setting stricter boundaries for the values of the parameters or starting with better initial estimates for *u*, the growth rate *r* and evolutionary speed *σ*.

The model is a first attempt to forecast the patient’s future tumor trajectory. The success at forecasting was strong when the first three measurements were used, at least for patient’s that did not show pseudo-progression. The model also invites computer simulated “i-trials” [35], by modelling how a patient’s tumor burden would have responded to different therapeutic strategies such as adaptive therapy, dose amplification, or other strategies based on (leader-follower) games [49, 58]. With the model presented here, stabilizing the tumors was not feasible, but it seems possible to slow down their growth in some cases, especially for tumors that start growing exponentially from the beginning (non-responder patients) or after a period of response to treatment. The reason for these impediments is that the growth rate of these tumors is too high to let them grow unrestricted for long periods of time. Therefore, in this type of cancer, treating continuously at maximum tolerable dose often yielded better predicted outcomes than adaptive therapy.

Combining immunotherapy with chemotherapy has become more attractive, yet to date, has provided mixed results in clinical trials and patient outcomes [14, 29]. Issues include risks of toxicity if the drugs are given together [44], and if administered sequentially how best to do so. Two hopeful lines of research are open: one involves modeling multi-drug adaptive therapy [57], and the other extending our present model to include two or more drugs and the ways that they might interact to influence each others’ efficacy [41, 13]. To be effective, such models would require more a priori information concerning the effect that the available treatments have on the growth of the tumors when drugs are administered alone or together.

We applied the same model to immunotherapy and to chemotherapy. The only difference being the parameterizations that emerged from fitting the respective patient data. As an additional difference, we treated the immunotherapy as either on or off, whereas the chemotherapy dosing existed on a continuum from 0 to 1 (maximum tolerable dose). While immune system models can explicitly consider the dynamics of other elements of the immune system [42, 38, 32, 18], when the patient data are just time series of tumor size, without any measures of immune infiltration or immune system dynamics, then the model we used here should suffice as an abstraction.

Our model showed a significant role for the rate of evolution of resistance. We allowed this parameter to be patient-specific. Its magnitude should reflect a patient’s overall tumor heterogeneity, mutation rates among the cancer cells, and epigenetic and phenotypic plasticity. For most patients the estimated value of this evolutionary speed term was greater than zero and permitted, in many cases, the rapid evolution of resistance. We conducted simulations exploring the consequences of therapeutically targeting the rate of evolution. This would have improved the final outcome for more than half of the patients, especially those who do not respond to treatment because of a too rapid evolution of resistance. How to target a cancer’s evolvability remains challenging both in theory and in practice. Hypomethylation appears to be one pathway by which cancer cells can explore their genome and create genetic instabilities associated with more rapid evolution [28]. This may be one explanation for the efficacy of DNA hypomethylating drugs [48].

Our model failed to fit the dynamics of tumors exhibiting pseudoprogression, i.e., tumors that may keep growing for a relatively long period of time after the start of treatment and then start responding to treatment. This poses a challenge to modelling and fitting tumor dynamics in general. If the tumor measurements showing pseudoprogression (via radiographic imaging) are an accurate reflection of the actual tumor burden, then introducing time lags into the model may account for such behavior. Alternatively, pseudoprogression may not reflect actual tumor burden but inflammation associated with the killing of cancer cells. Such inflammation may be indistinguishable from live tumor on imaging [19, 8]. The solution to this does not lie in the modelling but in methodologies and biomarkers for accurately measuring tumor volumes. Applying ours or any model as a decision tool requires a tight iterative approach with the timing and accuracy of a patient’s tumor burden and characteristics.

## Acknowledgements

The authors thank F. Hoffmann-La Roche Ltd. for sharing the raw data through the platform “Clinical Study Data Request” (CSDR, www.ClinicalStudyDataRequest.com) as part of the original study (REF). This research was supported by European Union’s Horizon 2020 research and innovation programs under the Marie Skłodowska-Curie grants 690817 and 955708 and the Dutch National Foundation projects ENWPR.020.006 and OCENW.KLEIN.277.

## Author contributions

Conceptualization, all; methodology, V.A.M., M.S., J.S.B., R.C., and K.S.; software, V.A.M. (all codes can be found on https://github.com/yo9299/NSCLC); validation, V.A.M and M.S.; formal analysis, V.A.M., M.S., and K.S.; investigation, all; resources, N.G.L. and J.N.K.; writing, original draft preparation, M.S., V.A.M., and K.S.; writing, review and editing, all; visualization, V.A.M. and M.S.; supervision, F.T., J.S.B., R.C., J.N.K. and K.S.; funding acquisition, J.N.K. and K.S. All authors have read and agreed to the published version of the manuscript.

## Supplementary material

